# Response of whitefly to the wild tomato *Solanum habrochaites*

**DOI:** 10.1101/2021.02.26.432993

**Authors:** Fengqi Li, Youssef Dewer, Jiahui Tian, Du Li, Cheng Qu, Zhen Yang, Chen Luo

**Affiliations:** Beijing Key Laboratory of Environment Friendly Management on Fruit Diseases and Pests in North China, Institute of Plant and Environment Protection, Beijing Academy of Agriculture and Forestry Sciences, Beijing 100097, China; Bioassay Research Department, Central Agricultural Pesticide Laboratory, Agricultural Research Center, Dokki, Giza 12618, Egypt; Tianjin University of Traditional Chinese Medicine, Tianjin 300000, China

**Keywords:** whitefly, *Solanum habrochaites*, transcriptome

## Abstract

The whitefly Bemisia tabaci (Gennadius) causes severe damage to cultivated tomato in many regions of the world through direct feeding and indirectly through transmission of plant viruses. Field observations show that B. tabaci is rarely infested the non-host plants such as the wild tomato Solanum habrochaites; however, the molecular mechanism involved in the recognition of wild plant odors is still unclear. In this study, we assessed the effects of S. habrochaites on the survival, fecundity, and egg hatchability of the Mediterranean (MED) species of B. tabaci. Expression and splicing of stress-response genes in whitefly exposed to S. habrochaites was analyzed using RNA-sequencing data and alternative splicing analysis. These results indicated that the S. habrochaites treatment can induce the expression of environmental stress genes in B. tabaci. This study may help us to a better understanding of the molecular mechanisms involved in the olfactory recognition of non-host volatiles particularly wild tomato relative. Furthermore, the findings of this study may provide excellent chances of finding a suitable antagonist of eco-friendly properties which can block the perception of chemosensory signals. Thereby, the feeding behavior and food preferences of B. tabaci can be manipulated and thus insect populations can eventually be controlled.

## 1. Introduction

The Whitefly *Bemisia tabaci* (Gennadius) is one of the most important pests that causes economic damage to crop plants (De et al., 1997). *B. tabaci* has a high oviposition rate and rapid population growth (Erdogan et al., 2008). The Middle East-Asia Minor1 (MEAM1) and Mediterranean (MED) subspecies are the most invasive *B. tabaci* whiteflies worldwide (Chu et al., 2006). In China, the MED cryptic species has now been replaced by MEAM1 in many regions (Hu et al., 2011).

*B. tabaci* is one of the main tomato pests that reduces tomato growth through direct feeding and indirectly through transmission tomato yellow leaf curl virus (TYLCV) (Su et al., 2016). *B. tabaci* is the only vector of TYLCV, so whitefly control is one of the key measures used to manage TYLCV (Wei et al., 2017). Chemical insecticides remain the main tools in the management of whitefly, but the rapid development of resistance to several chemical classes of insecticides has made it so difficult to control (Elbert et al., 2000). Therefore, there are urgent demands for safe alternative strategies in modern pest management, i.e. for controlling whitefly populations. One of these novel approaches is based on the fact that chemical communication via the olfactory system drives essential behaviors of insects (Ingham et al., 2020). Indeed, detecting appropriate chemical signals is essential for insects to find their mating partners, oviposition sites and food sources (Pelosi et al., 2017; Leal and Walter, 2013). Accordingly, the olfactory system of many insects has evolved to chemo-detectors of high sensitivity and accuracy, which allow the insect to detect plant volatile odors at extremely low concentrations and to discriminate a large variety of odor cues (Suh et al., 2014). Thus, understanding of the mechanisms and signals that are involved in non-host selection behavior of *B. tabaci* may lead to strategies that exploit the odorous alert signals to enhance crop resistance.

*B. tabaci* is attracted to cultivated tomato, but it is repelled by the wild tomato relative *S. habrochaites*. The repellent behavior response of *B. tabaci* exposed to *S. habroachaites* as non-host plant was found to be associated with the emission of 7-epizingiberene and R-curcumene compounds (Bleeker et al., 2009). This two terpene volatiles have shown strong repellency against *B. tabaci* and induced *S. habrochaites* to develop enhanced resistance to insect infestation (Freitas et al., 2002; Muigai et al., 2002). On the other hand, *B. tabaci* did not elicit behavioral responses to the two corresponding chiral isomers: α-zingiberene and S-curcumene extracted from ginger (Bleeker et al., 2011). Transformation of cultivated tomato with the gene encoding *S. habrochaites* 7-epizingiberene synthase has become strongly repellent to *B. tabaci* that reduced whitefly fecundity up to 87% (Bleeker et al., 2012). These studies showed that *S. habrochaites* can exert strong repellent and toxic stresses aganist *B. tabaci*, but it is unclear how it exerts its effects on the physiological and molecular levels.

In order to achieve the goal, it was necessary to study the expression-related genes that may be affected in the whitefly *B. tabaci* after exposure to the wild tomato relative *S. habrochaites*.

## 2. Materials and methods

### 2.1 Insects rearing and plant materials

The MED cryptic species of whitefly was reared on tomato plants (Xianke 8) under greenhouse conditions of 25 ± 2 °C, 65 ± 5% relative humidity (RH), and a 16:8 h (L:D) photoperiod. The tomato plant was placed into cage measuring (60 cm × 60 cm × 60 cm). The source of MED was collected from a greenhouse located in Beijing, China and identified as MED by mitochondrial cytochrome oxidase I (mtCOI) gene (Brown et al., 2005). One pair of newly emerged adults was used for colony initiation, and the MED source after six generations was used for subsequent experiments.

### 2.2 Identification of wild tomato volatiles

*S. habrochaites* were placed into the cages measuring (60 cm × 60 cm × 60 cm) and grown under greenhouse conditions of 25 ± 2°C, 65 ± 5% RH, and a 16:8 h (L:D) photoperiod. Six-week-old plants were used for volatile analysis. We placed leaves of a healthy tomato plant into a glass bottle (45 mL, cleman) and an SPME fiber 50/30-μm divinylbenzene/carboxen/polydimethylsiloxane (Supelco; Sigma-Aldrich, http://www.sigmaaldrich.com) was used for analyte extraction. GC-MS analysis was performed on an Agilent 7890 (Agilent Technologies, Tustin, CA, USA) gas chromatography and Agilent 5975 mass selective detector.. Samples were separated using both HP-5 and DB-WAX (both 30 m × 0.25 mm i.d., 0.25 mm film thickness, Agilent Technologies, Tustin, CA, USA). Helium was used as carrier gas at a flow rate of 1.7 mL/min, and the GC inlet was set in the split-less mode. The injector temperature was 220 °C. The optimization of method parameters of SPME and GC-MS were performed following to the method previously described (Li et al., 2018). R-curcumene has been identified using GC-MS techniques by comparing its retention time and mass spectrum with the same parameters from a reference compound. As the 7-epizingiberene is thermally unstable (Bleeker et al., 2011), therefore, the R-curcumene was used for conducting the behavioral and olfactory studies. The synthesis of R-curcumene and S-curcumene was performed according to the method previously reportedS (Song et al., 2012). The synthesized compounds were confirmed by using various spectral technique like mass spectrometry and NMR detection. The α-zingiberene was purchased from Sigma-Aldrich (St. Louis, MO, USA).

### 2.3 RNA sequencing and bioinformatics analysis

To evaluate the effect of *S. habrochaites* on whiteflies at the molecular level, RNA-sequencing was performed using MED females and males, which have been fed on tomato Xianke 8 and the wild tomato *S. habrochaites* LA2175 for 6 and 24 h, respectively. Sequencing libraries were constructed using strand-specific libraries. FastQC (Andrews et al., 2010) was used to assess the quality of the sequencing data while Fastp (Chen et al., 2018) was employed to remove adapters, contamination, and duplicate sequences. The sequenced clean data were mapped to the MED whitefly genome v1.0 (Xie et al., 2017) using HISAT2 (Kim et al., 2019) ftp://www.whiteflygenomics.org/pub/whitefly/MED/v1.0. FeatureCounts (Liao et al., 2014) was used to calculate the expression of each gene, edgeR (Roboison et al., 2010) was used to analyze the difference in gene expression (we used Log2 Fold Change>1, Adjusted P-value=0.05 as the threshold), rMATS (Shen et al., 2014) was used to analyze the alternative splicing, and Sashimi plot was used to draw the alternative splicing of genes. Finally, WGCNA (Langfelder and Horvath, 2008) was used to calculate the correlation between the gene expression level (TPM values) and the mortality phenotypes of whitefly.

## 3. Results and discussion

### 3.1 7-epizingiberene and R-curcumene are the main volatile components of *S. habrochaites*

We synthesized a high-purity R-curcumene and confirmed its identity by mass spectrometry and NMR (Supplementary Data 1). We found that 7-epizingiberene and R-curcumene are the main volatile compounds of *S. habrochaites* using SPME and GC-MS detection. GC-MS analysis revealed that these two volatile compounds were not detected in the volatile blends of common tomato plants (Fig. 1). It was also found that the peak areas ratio of 7-epizingiberene and R-curcumene in LA2175 was 2.37- and 4.27-fold that of LA2329, respectively (Fig. 1). The data showed that LA2175 leaves had a relatively higher content of terpene volatiles than LA2329. This is consistent with the biological analysis showing that LA2175 was more resistance to whiteflies. Therefore, we chose and employed LA2175 for subsequent whitefly high-throughput sequencing studies.

**Fig. 1:**
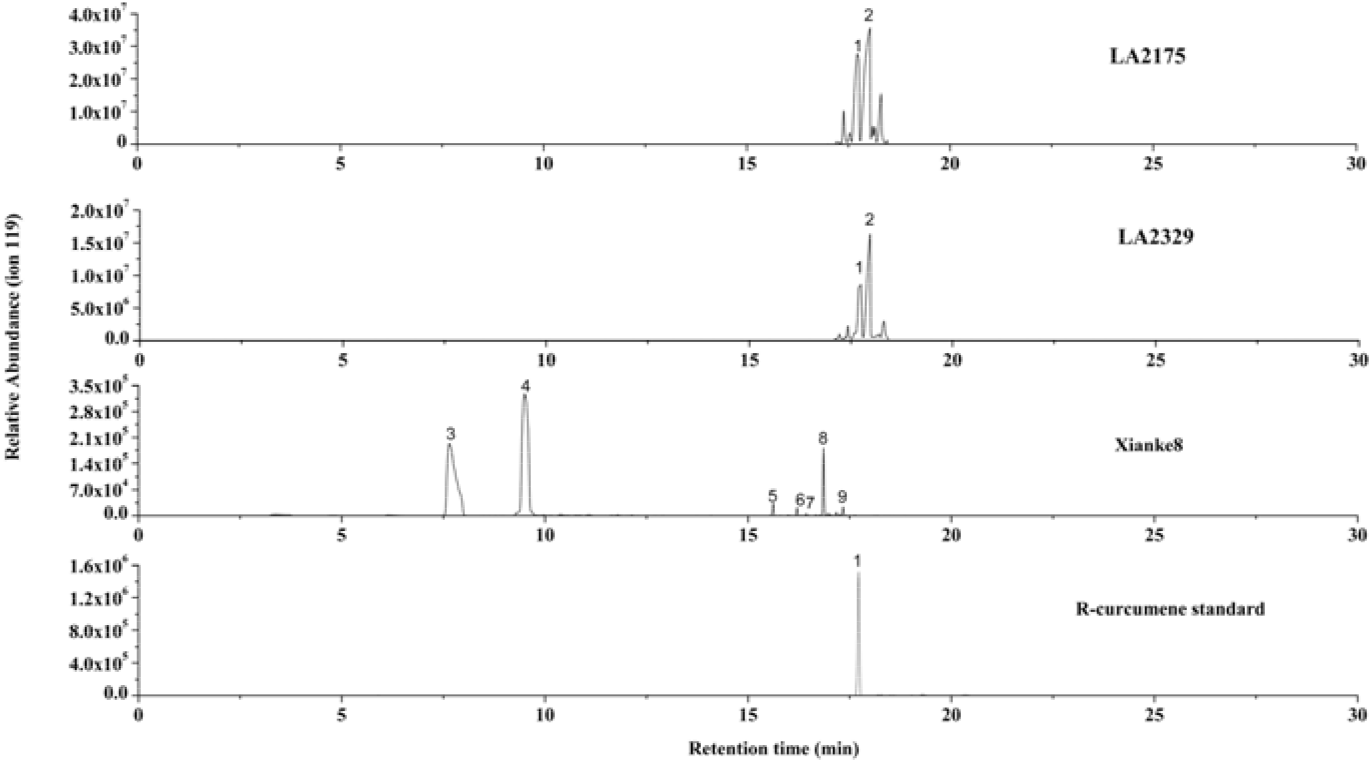
GC–MS analysis of the volatile profiles of wild tomato *Solanum habrochaites* LA2175, LA2329, and common tomato Xianke 8. 1. R-curcumene, 2. 7-epizingiberene, 3. *p*-cymene, 4. β-phellandrene, 5. elemene, 6. α-copaene, 7. unknow, 8.β-caryophyllene, 9. α-humulene.

### 3.2 Expression of whitefly-related genes after stress exposure of *S. habrochaites*

A total of 24 samples of male and female whitefly adults (including two time points and four treatments) were used for transcriptome sequencing (Fig. 2). About 21–24 million reads with an average length of 151 bp were aligned to the MED whitefly reference genome by HISAT2 (Table 1). About 78% of the reads mapped to unique loci (Table 1). After gene expression value analysis by featurecount and edgeR, we found four significantly upregulated genes in wild tomato 6-h-treated female adults (WF6) and wild tomato 24-h-treated female adults (WF24) (Fig. 2; Supplementary Data 2). They were BTA016109 (Multicopper oxidase), BTA027707 (Sulfotransferase), BTA001651 (OHCU decarboxylase), and BTA011604 (Heat shock protein). Wild tomato 6-h-treated male adults (WM6) and wild tomato 24-h-treated female adults (WM24) also included four genes that are both significantly upregulated (Fig. 2; Supplementary Data 2). They were BTA003886 (Heat shock protein), BTA016667 (Unknown), BTA005900 (serine threonine-protein kinase SBK1-like), and BTA011604 (Heat shock protein). One gene (BTA011604, Heat shock protein) was significantly upregulated together in these four treatments. There were two genes that were significantly downregulated in WF6 and WF24, BTA009028 (Unknown) and BTA013802 (Farnesyl-diphosphate farnesyltransferase). Eighteen genes were significantly downregulated in WM6 and WM24 (Fig. 2; Supplementary Data 2). Among these, BTA022129 is solute carrier family 3, member 1, and the other genes had unknown functions. These results indicated that the *S. habrochaites* treatment can induce the expression of environmental stress genes such as heat shock protein in whiteflies.

**Fig. 2:**
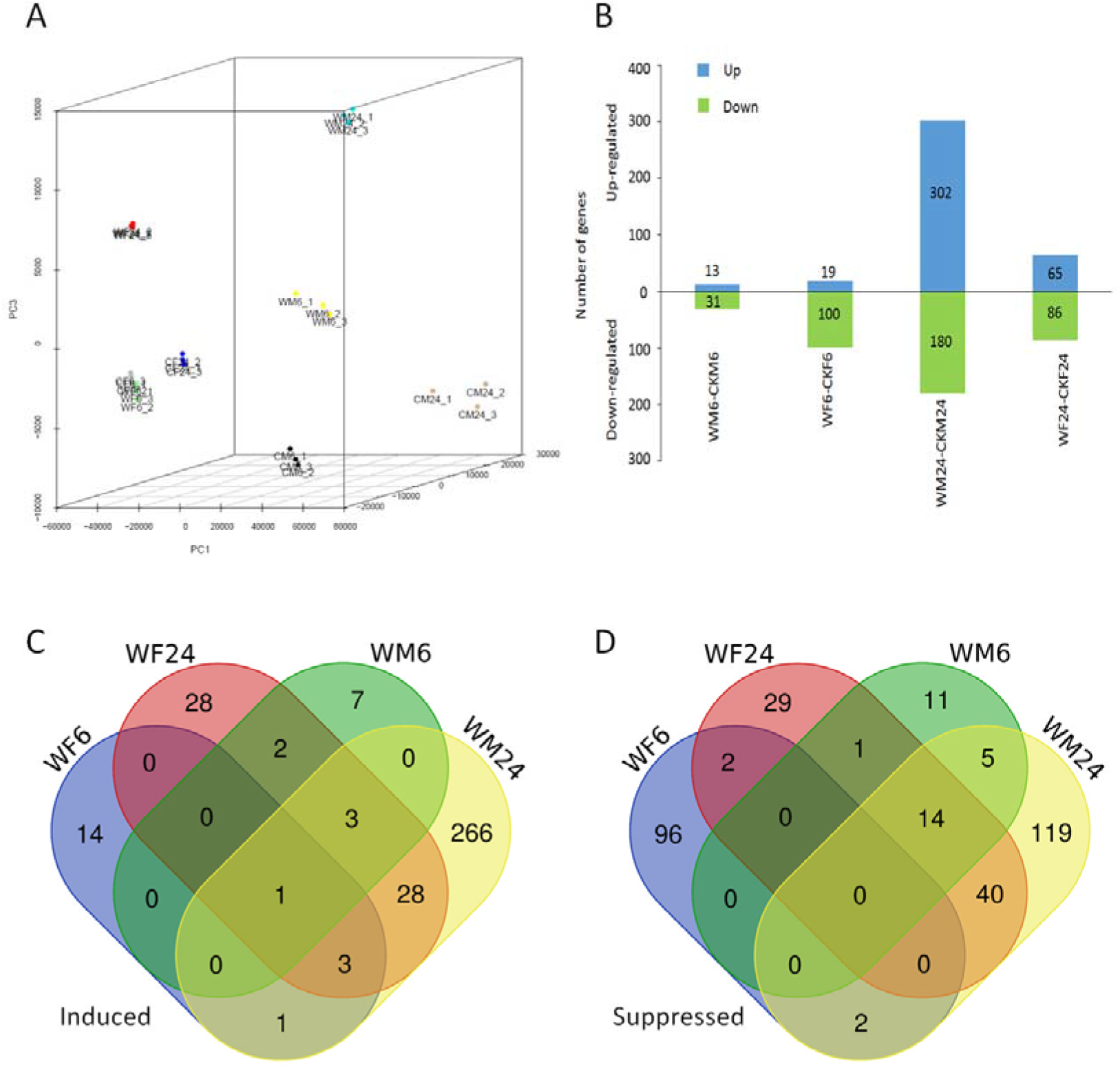
Analysis of the RNA-Seq data of whiteflies treated by common tomato and wild tomato. (A) Principal component analysis (PCA) plot showing clustering of RNA-Seq of whiteflies. (B) Bar graph showing the number of upregulated (upper bars) and downregulated (lower bars) genes in whitefly male and female adults. (C) Venn diagram showing the number of induced differently expressed genes unique to or common within the four conditions. (D) Venn diagram showing the number of suppressed differently expressed genes unique to or common within the four conditions. WF6: wild tomato (*Solanum habrochaites*) 6 hours treated female adults; WM6: wild tomato 6 hours treated male adults; WF24: wild tomato 24 hours treated female adults; WM24: wild tomato 24 hours treated male adults.

**Table 1.** Summary of the RNA-Seq Data. CKF6_1-3, whitefly females grown on common tomato for 6 h replicates 1–3. CKM6_1-3, whitefly males grown on common tomato for 6 h replicates 1–3. CKF24_1-3, whitefly females grown on common tomato for 24 h replicates 1–3. CKM24_1-3, whitefly males grown on common tomato for 24 h replicates 1–3. WF6_1-3, whitefly females grown on wild tomato LA2175 for 6 h replicates 1–3. WM6_1-3, whitefly females grown on wild tomato LA2175 for 6 h replicates 1–3. WF24_1-3, whitefly females grown on wild tomato LA2175 for 24 h replicates 1–3. WM24_1-3, whitefly males grown on wild tomato LA2175 for 24 h replicates 1–3.

|  | No. of paired-end reads | No. of mapped paired-end reads | Proportion of mapped paired-end reads |
| --- | --- | --- | --- |
| CKF6_1 | 22729871 | 19170373 | 84.34% |
| CKF6_2 | 24263436 | 20446798 | 84.27% |
| CKF6_3 | 24587112 | 20852330 | 84.81% |
| CKM6_1 | 22886414 | 18199276 | 79.52% |
| CKM6_2 | 22949620 | 18146265 | 79.07% |
| CKM6_3 | 22716792 | 18100740 | 79.68% |
| CKF24_1 | 22159172 | 18759955 | 84.66% |
| CKF24_2 | 22190660 | 18620183 | 83.91% |
| CKF24_3 | 23055250 | 19278800 | 83.62% |
| CKM24_1 | 22912092 | 18384663 | 80.24% |
| CKM24_2 | 22576192 | 17744887 | 78.60% |
| CKM24_3 | 22106168 | 17342289 | 78.45% |
| WF6_1 | 21695473 | 17994225 | 82.94% |
| WF6_2 | 25184704 | 21089671 | 83.74% |
| WF6_3 | 24590350 | 20636222 | 83.92% |
| WM6_1 | 23754024 | 19112488 | 80.46% |
| WM6_2 | 24277496 | 19358875 | 79.74% |
| WM6_3 | 22011538 | 17624639 | 80.07% |
| WF24_1 | 22934225 | 19340432 | 84.33% |
| WF24_2 | 22669251 | 18994565 | 83.79% |
| WF24_3 | 24285715 | 20487429 | 84.36% |
| WM24_1 | 23292401 | 19060172 | 81.83% |
| WM24_2 | 21426439 | 17460405 | 81.49% |
| WM24_3 | 24205186 | 19790160 | 81.76% |

Alternative splicing is often related to the response of the organism to a stressful environment (Sultan et al., 2008; Ling et al., 2015; Wang et al., 2008). We determined if whiteflies were stressed by exposure to *S. habrochaites* and whether genes were able to respond to the stress through alternative splicing. We performed alternative splicing analysis on differently treated samples (comparison of male and female samples treated with *S. habrochaites* and then treated with common tomato) through rmats. There were five types of alternative splicing, including skipped exon (SE), mutually exclusive exon (MXE), alternative to 5′splice site (A5SS), alternative to 3′ splice site (A3SS), and retained intron (RI). Among these, only SE and MXE had significant differences between the two treatments (FDR <0.05), and the SE type accounted for the main proportion (Table 2: Supplementary Data 3). GO enrichment analysis showed that genes with alternative splicing of the SE type were significantly enriched in three GO items including O-acetyltransferase activity, glycerol ether metabolic process, and ether metabolic process (Supplementary Data 4). The MXE-type alternative splicing genes were not significantly enriched.

**Table 2.** Genome-wide effects of *Solanum habrochaites* stress on alternative splicing of whitefly. WF6: wild tomato (*Solanum habrochaites*) 6-h-treated female adults; WM6: wild tomato 6-h-treated male adults; WF24: wild tomato 24-h-treated female adults; WM24: wild tomato 24-h-treated male adults. CKF6: common tomato 6-h-treated female adults; CKM6: common tomato 6-h-treated male adults; CKF24: common tomato 24-h-treated female adults; CKM24: common tomato 24-h-treated male adults.

| AS Events Summary | WF6 vs<br>CKF6 | WM6 vs<br>CKM6 | WF24 vs<br>CKF24 | WM24 vs<br>CKM24 |
| --- | --- | --- | --- | --- |
| SE | 7 | 13 | 11 | 14 |
| MXE | 0 | 1 | 2 | 5 |
| RI | 0 | 0 | 0 | 0 |
| A5SS | 0 | 0 | 0 | 0 |
| A3SS | 0 | 0 | 0 | 0 |

## Data availability conflict of interest

RNA-seq data have been deposited and are available in NCBI (BioProject ID is PRJNA622388).

