## Supplementary figures and images for "Response of whitefly to the wild tomato *Solanum habrochaites*"

### Supplementary Data 1

The result of mass spectrometry and NMR to confirm the identity of R-curcumene

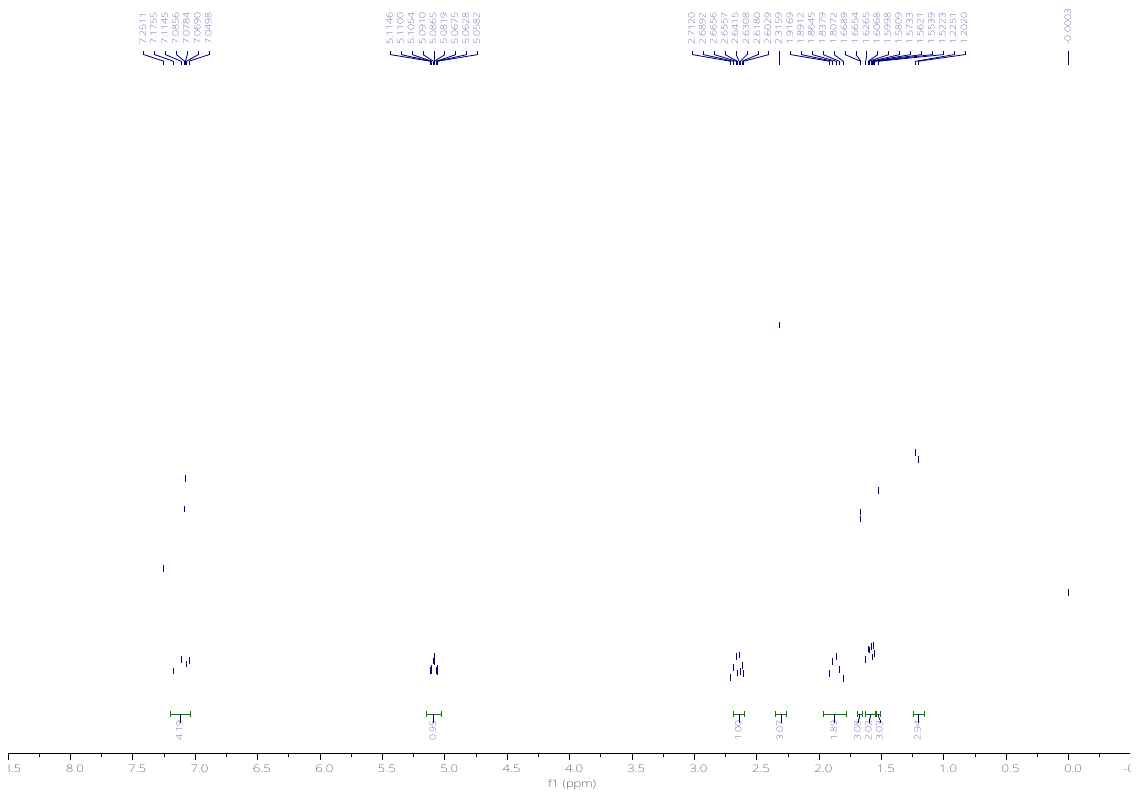

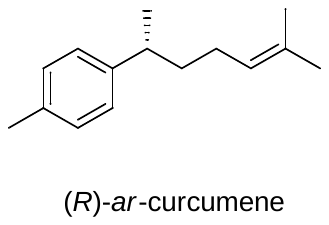

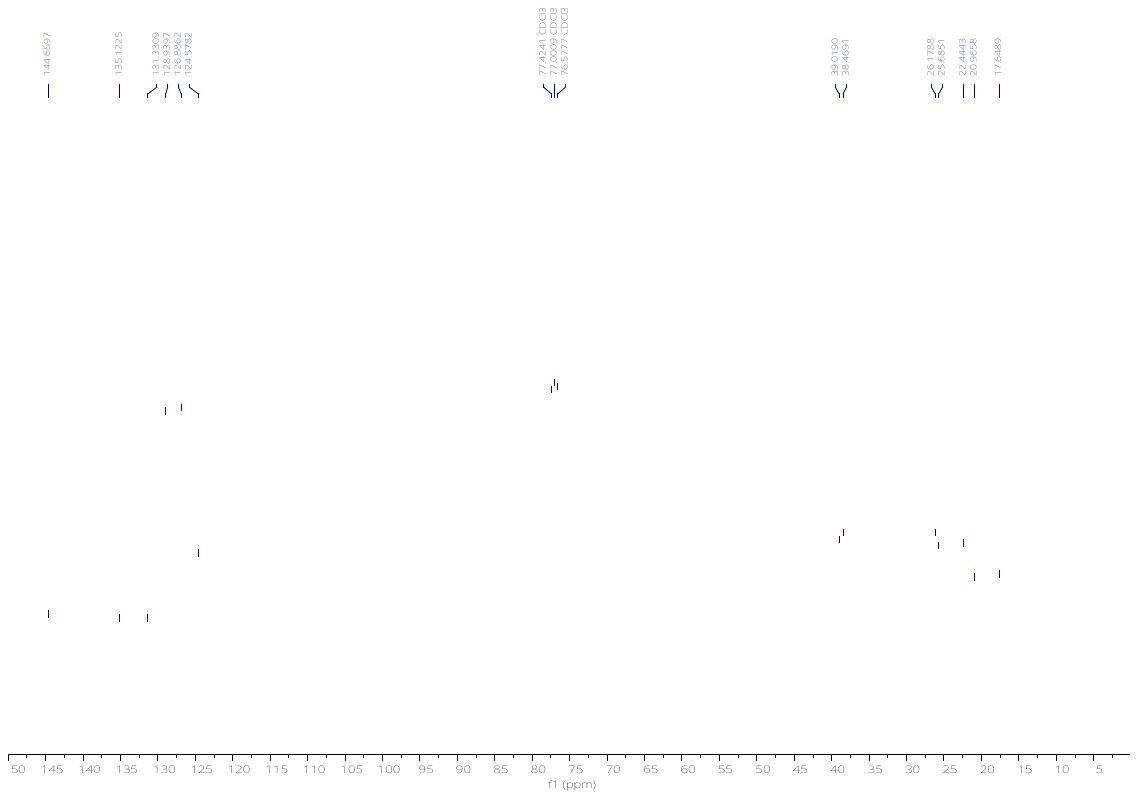

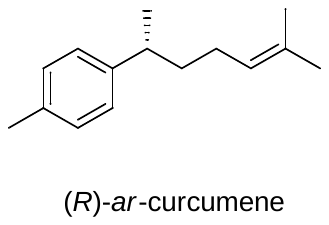

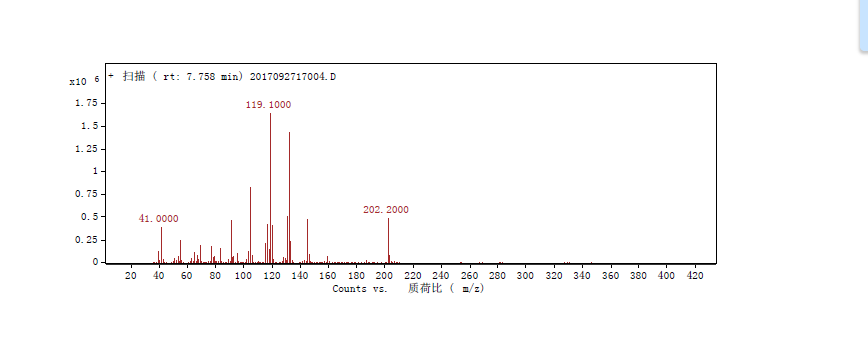
